# A dynamic link between respiration and arousal

**DOI:** 10.1101/2023.10.06.561178

**Authors:** Daniel S. Kluger, Joachim Gross, Christian Keitel

## Abstract

Viewing brain function through the lense of other physiological processes has critically added to our understanding of human cognition. Further advances though may need a closer look at the interactions between these physiological processes themselves. Here we characterise the interplay of the highly periodic, and metabolically vital respiratory process and fluctuations in arousal neuromodulation, a process classically seen as non-periodic. In data of three experiments (N = 56 / 27 / 25 women and men) we tested for covariations in respiratory and pupil size (arousal) dynamics. After substantiating a robust coupling in the largest dataset, we further show that coupling strength decreases during task performance compared with rest, and that it mirrors a decreased respiratory rate when participants take deeper breaths. Taken together, these findings suggest a stronger link between respiratory and arousal processes than previously thought. Moreover, these links imply a stronger coupling during periods of rest, and the effect of respiratory rate on the coupling suggests a driving role. As a consequence, studying the role of neuromodulatory arousal on cortical function may also need to consider respiratory influences.

**Significance statement:** We characterise the interplay of the respiratory rhythm and pupil diameter dynamics as a well-known proxy for arousal. Although we consistently find respiratory modulation of pupillary changes, they were most pronounced during periods of rest (compared to during task performance) and dependent on respiratory rate (deep vs. normal breathing).

## Introduction

An increasing number of findings embed rhythmic brain activity in a flurry of periodic physiological processes. Cardiac (Al et al., 2020; Galvez-Pol et al., 2020), gastric (Rebollo et al., 2021, 2018), respiratory (Kluger et al., 2021; 2023; Park et al., 2020), and arousal neuromodulation (Schneider et al., 2016; Groot et al., 2021) have been shown to impact cortical rhythms linked to human cognition. Fully grasping the role of physiological dynamics of the periphery for human cognition will also require understanding how these interact with each other (Kluger et al., 2024). Here, we investigate the potential co-variations of endogenous dynamics of respiration and pupil-linked arousal.

The breathing rhythm has attracted particular interest because it can be voluntarily controlled (Allen et al., 2023; Brændholt et al., 2023). Breathing arises from respiratory pattern generators in the preBötzinger complex of the brain stem (Del Negro et al., 2018). Efferent signals project to limbic and sensorimotor cortical areas (Yang and Feldman, 2018) via suprapontine nuclei and the central medial thalamus. Cortical activity in turn evokes changes in the primary respiratory network, e.g., to initiate transitions between brain states like heightened arousal during a panic attack. A recent systematic review by Schaefer and colleagues (2023) showed that respiratory coupling to arousal dynamics, as measured by pupillometry, remains critically understudied.

Arousal describes a global physiological preparedness to process and respond to sensory stimulation (Aston-Jones and Cohen, 2005). Physiologically, arousal levels are under the control of brainstem and basal forebrain nuclei: The locus coeruleus (LC) and the nucleus basalis of Meynert respectively control the release of norepinephrine and acetylcholine to widespread cortical regions (Hasselmo, 1995; Lee and Dan, 2012; Steriade, 1996), which regulate momentaneous wakefulness and transitions between behavioural states (Harris and Thiele, 2011; McGinley et al., 2015b; Schwalm and Rosales Jubal, 2017): Coincidentally, release of these neurotransmitters also affects pupil size, making pupillometry an effective readout of arousal (McGinley et al., 2015a; Reimer et al., 2014; Vinck et al., 2015). Variations in pupil-linked arousal have been shown to influence cortical activity (Pfeffer et al., 2022; Radetz and Siegel, 2022) and cognitive function (Waschke et al., 2019; Kosciessa et al., 2021).

During prolonged periods of low arousal, a resting pupillary rhythm emerges, the so-called Hippus (Mathôt, 2018). The Hippus has therefore been used as an indicator of declining alertness or increasing drowsiness (McLaren et al. 1992). Interestingly, its typical peak frequency (0.2 Hz; Bouma and Baghuis, 1971) roughly coincides with the average pace of the breathing rate.

The relative neighbourhood of respiration- and arousal controlling brainstem structures, as well as the existence of concrete synaptic connections from the preBötzinger complex to LC suggests that both functions interlink (Ohtsuka et al., 1988). In fact, interrupting the connection between both cores in the rodent brain led to chronic hypoarousal and “lethargic” behaviour (Yackle et al., 2017). Although a recent review failed to find conclusive evidence for links between respiratory and pupil dynamics in humans (Schaefer et al., 2023), we hypothesise that their anatomical and functional interactions can be reflected in the respiratory modulation of pupil size, specifically the Hippus. Due to the periodic nature of both signals, a co-variation may express as phase coupling (Gross et al., 2021).

Considering this, we tested whether an emerging Hippus would indicate a stronger coupling of arousal to the breathing rhythm (Nakamura et al., 2019; Ohtsuka et al., 1988) in N = 56 resting, healthy volunteers. In a second dataset (N = 27), we tested how the coupling changed when participants engaged in voluntary deep (vs normal) breathing. Finally, for a subset of participants of the first study (N = 25), we compared spectral characteristics of respiratory and pupil-linked arousal dynamics between rest and during task performance and tested whether a potential coupling would depend on their behavioural state. We show evidence for a coupling of respiratory and pupillary dynamics during rest. Coupling characteristics changed with respiratory rates and coupling strength decreased during task engagement.

## Materials and Methods

### Participants and procedures

Respiratory and pupil data of N = 56 right-handed volunteers (31 female, age 25.3 ± 3.1 years [M ± SD]) were originally recorded for MEG studies published elsewhere (Kluger and Gross, 2020); Kluger et al., 2023; Pfeffer et al., 2022). All participants reported having no respiratory or neurological disease and gave written informed consent prior to all experimental procedures. The studies were approved by the local ethics committee of the University of Münster (approval numbers 2018–068f-S and 2021-785-f-S). During all procedures, participants were seated upright while we simultaneously recorded respiration with a respiration belt transducer around the chest (BIOPAC Systems, Inc, Goleta, USA) and pupil area of the right eye with an EyeLink 1000 plus eye tracker (SR Research). Both, respiration and eye tracking data, were sampled at 600 Hz.

All N = 56 participants completed a 5 min resting-state recording during which they were to keep their eyes on a fixation cross centred on a projector screen placed in front of them. A subset of n = 27 participants (12 women, age 25.0 ± 2.8 years) took part in a second study (see Kluger and Gross, 2020) in which they were instructed to breathe either naturally or voluntarily deeply through the nose for two separate 5 min runs.

A second, independent sample of n = 25 participants (13 female, age 25.5 ± 2.7 years) took part in a third study (see Kluger et al., 2021) in which they performed a simple visual detection task: Gabor patches were presented for 50 ms at near-threshold contrast either to the left or to the right of a central fixation cross displayed on a projector screen in front of them. After a short delay of 500 ms, participants were to report via button press whether they had seen the target on the left, the right, or no target at all. For comparison with the resting condition, we report results from the last of six task runs (120 trials, mean duration 446 ± 34 sec).

### Data preprocessing

Eye tracking and respiration time series of all three datasets used in analyses underwent largely similar preprocessing steps. Analyses made use of the Matlab toolbox fieldtrip (Oostenveld et al., 2011) in combination with custom-written code. All analysis scripts can be found on the Open Science Framework (https://osf.io/ysb3g/?view_only=f592983532b945ea9cd6000d8ffa6037).

Pupil area traces were converted to pupil diameter to linearize our measure of pupil size. Using routines available from https://github.com/anne-urai/pupil_preprocessing_tutorial (also see Urai et al., 2017), blinks were identified by an automatic (and visually validated) procedure and linearly interpolated. We slightly modified the original procedure in that blinks were detected in a first pass with a standard criterion of *z* = 3 SD. Then, blink-interpolated pupil time series were subjected to the procedure again using a relaxed criterion of *z* = 6 SD to capture remaining artifacts. Next, canonical responses to blinks were estimated and removed from pupil time series (Hoeks and Levelt, 1993; Knapen et al., 2016; Wierda et al., 2012). To that end, pupil time series were band-pass filtered (pass band: 0.01–10 Hz, second-order Butterworth, forward-reverse two-pass). Due to its robustness against artifacts, Respiration data underwent the same filtering procedure but no other preprocessing. For analysis, all time series were converted to z scores using a robust procedure (MATLAB function ‘normalize’ with options set to ‘zscore’ and ‘robust’).

### Data analysis

#### Spectral analysis

Power spectra of full-length recordings for all three datasets were obtained by subjecting pupil- and respiration time series to spectral decomposition by means of the multi-taper method as implemented in fieldtrip. Spectral smoothing was set to 0.05 Hz. All time series were zero-padded to a length of 600 sec to match the spectral resolution irrespective of the original length of the recordings. We used a logarithmically spaced frequency axis, with a frequency range of interest from 0.065 to 6Hz, avoiding the influence of filter artifacts (see Data preprocessing). As a final step, all power spectra were converted to dB scale by taking the decadic logarithm and multiplying by 10.

For pairwise comparisons between power spectra, we generally used the cluster-based permutation approach based on dependent-samples T-tests, and clustering results along the frequency dimension (*n* = 5000 permutations). Generally, cluster-corrected p-values had to satisfy a threshold of α = .05.

#### Respiration-pupil coupling – coherence

Coupling between respiration and pupil time series was quantified using the phase-based magnitude-squared coherence metric (Carter et al., 1973). To this end, we used the same approach as described above, but retained complex fourier spectra, and subjected these to a coherence analysis as implemented in fieldtrip (ft_connectivityanalysis, option ‘method’ set to ‘coh’, and ‘complex’ to ‘complex’). Additionally, this analysis was run on a combination of the original respiratory time series and a surrogate pupil time series. Surrogates were generated based on the spectral characteristics of the original pupil time series, retaining power-but scrambling phase information using the iterative amplitude-adjusted Fourier Transform approach (IAAFT; Schreiber and Schmitz, 1996; also see Kluger et al., 2023). The logarithmized magnitude-squared coherence spectra for respiration coupling with original and surrogate pupil data were then subjected to the cluster-based statistics as described for the power spectra above. For the visualisation of the distribution of individual data points in Fig. 1E (as well as 2B, and insets in Fig. 3A-C) we used the raincloud plot functionality for MATLAB (Allen et al., 2019; as implemented here: https://github.com/RainCloudPlots/RainCloudPlots/tree/master/tutorial_matlab).

**Figure 1.**
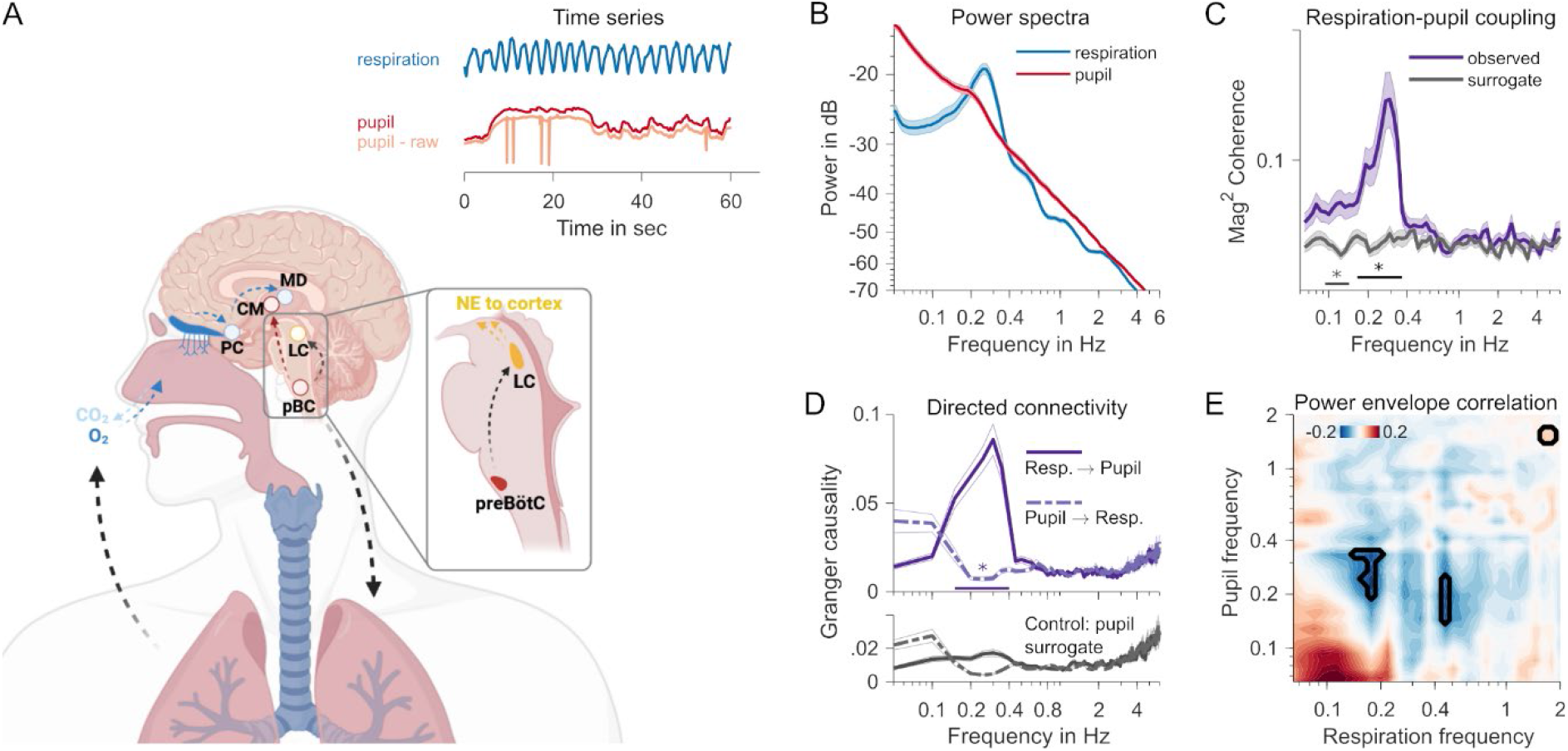
**A** Neuroanatomy of respiratory and pupillary rhythms. Respiratory pacemaker cells in the preBötzinger complex (pBC) project to locus coeruleus (LC), which controls global release of noradrenaline (NE), and to centro-medial thalamus (CM). In a feedback loop, the incoming airstream of each breath triggers receptors reaching from the epithelium into the nasal cavity, propagating breathing-locked activity to the piriform cortex (PC) and mediodorsal thalamus (MD). Inset time series show representative measurements of typical respiratory and pupil dynamics over the course of 60 seconds. The blue respiration trace shows the clear rhythmicity of breathing. A rhythmic component is not immediately visible in the red pupil trace. The raw unprocessed pupil trace, also depicted in orange, still contains the typical blinks (spikes) that have been interpolated for analysis (see Methods for details). **B** Spectral composition (power spectra) of 5-min resting pupil (red) and respiration (blue) time series. Shaded area shows SEM. **C** Mag-squared coherence spectrum between pupil and respiration time series. Observed coherence (purple) plotted against coherence computed on surrogate pupil time series (grey). Black straight line shows frequency range with significant differences (p < .05, cluster-based permutation test). **D** Directed connectivity quantified with spectrally resolved Granger causality. Top panel shows respiration-to-pupil (solid line) and pupil-to-respiration (dashed line) influences. Straight line at the bottom with asterisk indicates frequency range where connectivity differs (cluster-based permutation statistic, *p* < .05). Bottom panel shows connectivity spectra obtained from a control analysis using the surrogate pupil time series. **E** Power envelope correlations (Pearson’s correlation coefficient) between respiration and pupil time series. Marked areas denote significant differences at *p* < .05 (uncorrected; but relaxed cluster criterion applied - see Methods).

**Figure 2.**
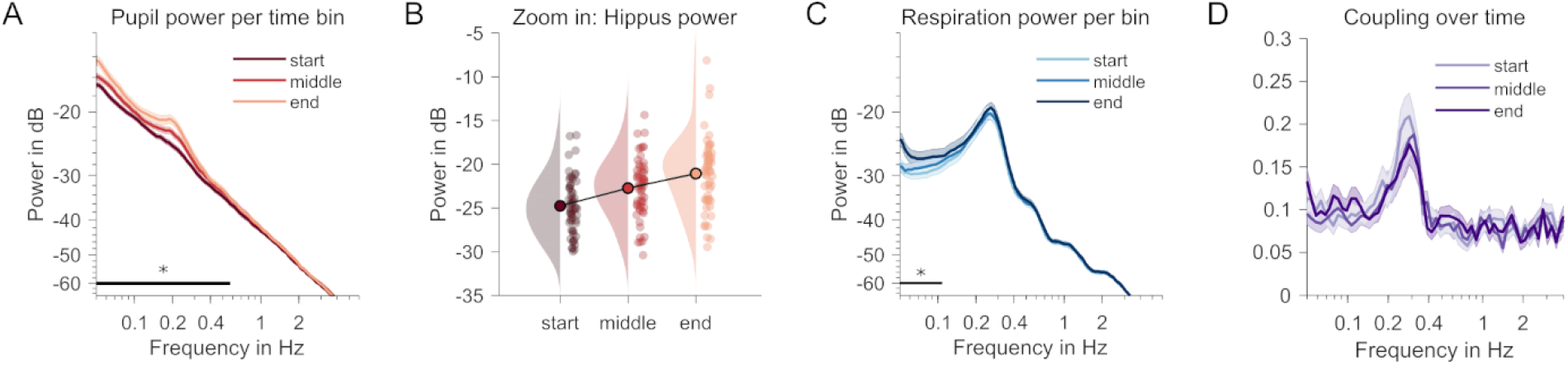
Changes over time **A** Pupil power spectra for three consecutive 2-min time windows. Darker colour = later time window. Black straight line shows frequency range with significant changes between time windows (p < .05, cluster-based permutation test). **B** Individual power values per time window at the Hippus peak frequency (0.19 Hz) in A. **C** Respiration power spectra, otherwise same as A. **D** Coupling (magnitude-squared coherence spectra) between pupil and respiration, otherwise same as A and C. No significant monotonic changes observed between time windows.

**Figure 3.**
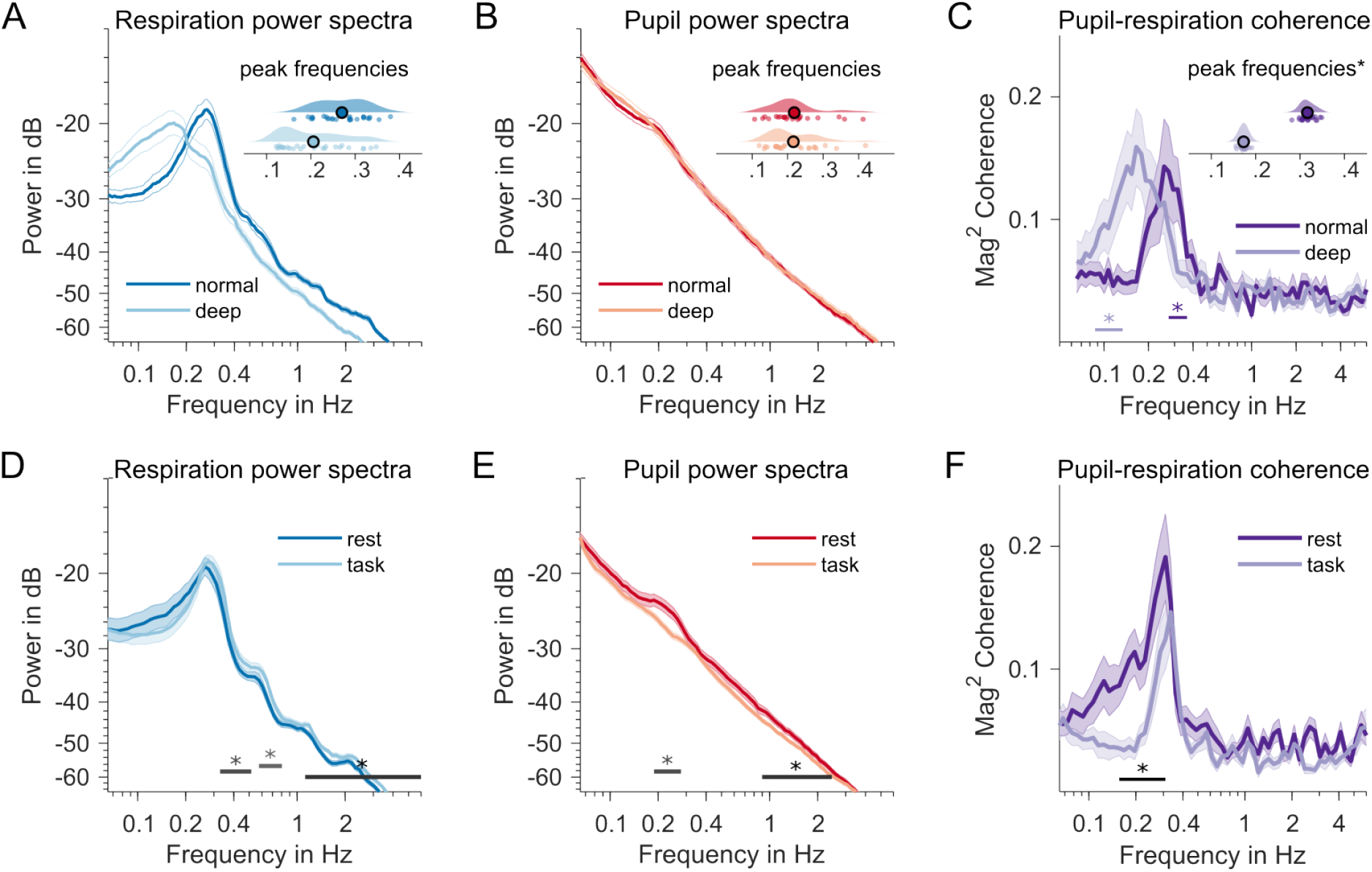
**A** Contrasted normal vs deep breathing conditions (dark vs light colours in A-C). Marked decrease in respiration frequency. Inset raincloud plot shows individual distributions of peak frequencies (in Hz) in both conditions. Offset along y-axis non-informative and for visualisation only. **B** Same as in A but for pupillary dynamics. No significant differences in Hippus peak frequencies between normal and deep breathing. **C** Individual coupling peak frequencies (* = variance-corrected jackknife estimates) are markedly reduced during deep breathing. Straight lines below spectra signify systematic coherence differences when compared with surrogate data (not shown). **D** Contrasted rest vs task conditions (different sample than A-C). Respiration power spectrum during rest (dark blue) and task performance (light blue). Straight lines with asterisks at the bottom of D, E and F indicate frequency ranges with significant changes between both behavioural states (p < .05, pairwise t-statistic, cluster-based permutation test). **E** Same for power spectra of pupil time series, dark red = rest, light red = task. **F** Comparison of respiration-pupil coupling, measured as magnitude-squared coherence. Dark purple = rest, light purple = task condition.

Identical approaches were taken when testing respiration-pupil coupling (coherence) in rest-vs-task and normal-vs-deep breathing datasets.

#### Respiration-pupil coupling - directed connectivity

We also tested whether the coupling between respiration and pupil showed a specific directionality that may suggest a causal influence in the resting-state dataset (N = 56). Again, we extracted complex Fourier spectra from the pre-processed respiration- and pupil time series by means of the multi-taper method. Note that due to the specific requirements of this analysis in fieldtrip, frequencies were linearly spaced, ranging from 0 to 6 Hz in 0.05 Hz steps, and spectral smoothing was set to 0.1 Hz. This analysis was also applied to a surrogate pupil time series, generated with IAAFT as explained above. Complex spectra were then used to estimate Granger causality (Bressler & Seth, 2011), using the nonparametric method as implemented in the fieldtrip function ft_connectivityanalysis (‘method’ set to ‘granger’, otherwise using default settings) for each participant individually. This analysis yielded spectra that indicated the directional connectivity respiration → pupil and vice versa, and similarly for the pupil surrogate time series. These were subjected to two statistical analyses, first, a cluster-based permutation test of spectrally resolved Granger causality between the two directions respiration → pupil and vice versa, and secondly, similar pairwise contrasts between the original data and the Granger causality estimates based on pupil surrogate data (e.g., respiration → pupil vs respiration → pupil surrogate). We note that the results of this analysis must be interpreted with caution as Bastos & Schoffelen (2016) have pointed out that directed connectivity measures can underlie a signal-to-noise ratio (SNR) issue that may lead to spurious results. Our measures of respiration and pupil dynamics naturally differ in SNR (see Figure 1B) and their respective measurements entail distinct noise characteristics.

#### Respiration-pupil coupling - power correlation

In the resting-state only dataset (N = 56), we also tested for potential covariations in power of respiration and pupil dynamics over the course of the 300 sec of the recording. To this end, time series were subjected to a spectro-temporal decomposition using the multitaper approach as implemented in fieldtrip (function ‘ft_freqanalysis’, with ‘method’ set to ‘mtmconvol’). Again, data were zero-padded to 600sec, and spectral smoothing was set to 0.2 Hz. For 36 frequencies, spanning a range from 0.065 to 2 Hz logarithmically, spectral representations were computed for frequency-dependent time windows of nine cycles each, and with a temporal step size that had consecutive windows overlap by 50%. The following time-frequency representations (power envelopes) for respiration data were correlated (Pearson’s correlation coefficient) with power envelopes of the pupil data, after dB-scaling them. Correlation coefficients were transformed to Fisher’s *z* before subjecting to statistical testing. As a contrast, we correlated original respiration power envelopes with surrogate pupil power envelopes, again derived from IAAFT (Schreiber and Schmitz, 1996; see Kluger et al., 2023; also see section ‘Respiration-pupil coupling - coherence’).

Initially, *z*-transformed original correlation coefficients were then compared against the surrogate data by means of cluster-based permutation testing using pairwise comparisons (‘ft_statfun_depsamplesT’) in the two-dimensional frequency-frequency plane (resulting from correlating respiration power envelopes of every frequency with pupil envelopes of every frequency, and vice versa). Prior to the testing a blurring filter was applied to individual frequency-frequency planes (original and surrogate correlations) to reduce the influence of small interindividual differences in frequencies (MATLAB function ‘imgaussfilt’, default settings). This analysis did not produce any clusters that survived correction for multiple comparisons. We then applied a relaxed cluster thresholding to a map obtained by applying paired t-tests to data at each frequency-frequency pair. In this map a cluster was only considered significant when consisting of at least five adjacent frequency-frequency correlations below the uncorrected α threshold. Correlation values (not *z*-transformed) were averaged across frequency-frequency correlations of the overall largest cluster (and displayed in Fig. 1E). Note that this relaxed criterion increases the chance of false positives.

#### Time-window analyses

For analysing changes over time in the resting-state only dataset (N = 56), preprocessed time series (all 300 sec in length), were re-epoched into three overlapping time windows of 150 sec each, with 50% overlap between time windows. Windowed data underwent the same spectral decomposition approach as described in section ‘Spectral analysis’. To test for monotonous trends, increases or decreases, in pupil and respiratory power, as well as in pupil-respiratory phase coupling (by means of magnitude-squared coherence) across the time windows, we used the cluster-based permutation approach based on dependent-samples regression, as implemented in fieldtrip (‘ft_statfun_ depsamplesregrT’).

#### Peak frequency shifts in normal vs deep breathing comparisons

When analysing the normal-vs-deep breathing dataset it became clear that the major effect in respiratory, pupil and coupling dynamics was a change in peak frequencies rather than spectral power (or magnitude-squared coherence when looking at coupling). To capture this effect, we refrained from the general approach to testing spectral differences described above and instead extracted individual peak frequencies as follows: First, we computed the spectra of the first-order gradient respiration and pupil time series (same parameters as reported in section ‘Spectral Analysis’). This effectively removes the 1/f component from resulting power spectra. Then, we increased the resolution of individual spectra by a factor of 100 using spline interpolation (MATLAB function ‘interp1’, option ‘spline’). This allowed capturing finer differences in the exact individual peak frequencies. These were extracted by means of the ‘findpeaks’ function (MATLAB), applied to individual spectra in the frequency range between 0.1 and 1 Hz. A similar approach was used to extract peak frequencies from coherence spectra (obtained as described in section ‘Respiration-pupil coupling - coherence’). Given their lower signal-to-noise ratio, we detected peak frequencies in leave-one-out sub-averages instead, and corrected resulting estimates for deflated variance following Smulders (2010). Outlier estimates (individual peak frequencies, 3 standard deviations from the mean) have been excluded from depictions in **Fig 3A-C** but have been left in the data for statistical comparisons. To reduce their influence on the overall outcome we used non-parametric Wilcoxon signed-rank statistics to test differences in peak frequencies for respiration and pupil power spectra, as well as respiration-pupil coherence spectra.

## Results

First, we analysed continuous 5-min long measurements of pupil diameter and respiratory dynamics from N = 56 resting volunteers who took part in a series of MEG studies. Analyses that investigated the relationships of MEG-recorded cortical activity with either pupil or respiratory dynamics have been published elsewhere (Kluger et al., 2023, 2021; Pfeffer et al., 2022). Here, we focus on the link between respiratory dynamics and changes in pupil size as an indicator of arousal neuromodulation.

### Coupling between respiration and pupil-linked arousal at rest

Separate spectral analyses of pupil- and respiration time series showed that respiration fluctuated periodically at the typical breathing rate of about 0.25 cycles per second (Fig. 1B). Further, smaller spectral peaks can be observed at harmonic frequencies due to the non-sinusoidal waveform of the respiratory fluctuations. Although less pronounced when compared with the respiration spectrum, pupil dynamics also contained a distinct periodicity that peaked at 0.19 Hz. This is consistent with the previously described Hippus, a resting rhythm of the pupil (Bouma and Baghuis, 1971; Pomè et al., 2020).

Next, we tested for the coupling of both signals by means of spectral cross-coherence. Individual coherence values, quantified as the logarithm of the amplitude of the coherence spectrum were subjected to a cluster-based permutation procedure that tested against surrogate coherence spectra based on reconstructed pupil time series with identical power-but scrambled phase spectra (IAAFT approach; Schreiber and Schmitz, 1996; also see Kluger et al., 2023). This analysis returned a cluster between 0.17 and 0.37 Hz in which coherence in the original data systematically exceeded the coherence expected by chance (*T*_*sum*_ = 41.57, *p*_*cbpt*_ < .001; Fig. 1C). This cluster contained the peak coherence at 0.29 Hz. A smaller adjacent cluster was found between 0.09 and 0.14 Hz (*T*_*sum*_ = 19.03, *p*_*cbpt*_ = .002; Fig. 1C). This extra cluster could be due to a subharmonic process stemming from the periodic but non-sinusoidal waveforms of the underlying physiological signals. We also note that smaller differences between exact peak frequencies (respiration or Hippus compared with coupling) should not be overinterpreted as a mismatch because the coherence estimate is based on two inherently noisy signals.

Testing for directionality in the respiration-pupil coupling, using a non-parametric spectrally-resolved Granger causality metric, showed an increase in the 0.15 to 0.4 Hz frequency range that was specific to the respiration-to-pupil direction (compared with pupil-to-respiration: *Tsum* = 31.51, *pcbpt* = .002; see Figure 1D, top panel) suggesting that the respiratory rhythm (Granger-)causes rhythmic pupil fluctuations in the Hippus frequency range. This effect was exclusive to using the original pupil data when compared with the same directional connectivity (respiration-to-pupil) but calculated using surrogate IAAFT-generated pupil time series (*Tsum* = 39.99, *pcbpt* = .002; compare Figure 1D top and bottom panel). The latter result argues against an SNR issue sensu Bastos & Schoffelen (2016) as the pupil surrogate time series shares the SNR and noise characteristics of the original pupil trace.

We also tested whether arousal- and respiratory signals covary in magnitude by subjecting wavelet-decomposed time series, i.e., power envelopes, to a frequency-frequency correlation analysis. Again, this was tested against correlations with a surrogate pupil time series generated by means of IAAFT (Fig. 1E). This analysis produced a cluster of negative correlation that coincided with the peak frequencies of respective power spectra, peaking at 0.18 Hz (respiration) and 0.28 Hz for the pupil (cluster peak Pearson’s *r(54)* = -0.19, *p*_*uncorr*_ < .05 for a minimum of five adjacent pixels in the frequency-frequency plane; a cluster-based permutation test did not return any significant clusters). A similar smaller cluster at a higher respiration frequency mirrored the effect and was likely due to the strong harmonics in the spectral composition of the respiration time series (see Fig. 1B). As the peak power of the respiratory rhythm is a measure of the depth of breathing, this suggests that shallower breathing coincides with a stronger pupillary Hippus. An interpretation of this negative association may be that, during rest, episodes of lower metabolic requirements reduce respiratory depth, yet increase the coupling of arousal to respiration producing a stronger Hippus.

### Hippus increases during rest

To explore changes in the coupling of respiratory and pupil dynamics over time, we split the 5 min resting state recordings into consecutive overlapping segments of 150 sec (3 segments, 50% overlap). Power spectra of each segment were then submitted to a regression analysis using cluster-based permutation testing to identify frequency ranges with monotonous changes across segments. These analyses identified a cluster centred on the Hippus frequency range for pupil dynamics (*T*_*sum*_ = 174.52, *p* = .001) and ranging from 0.05 to 0.58 Hz (Fig. 2A & B). We also found an increase in power in the frequency ranges between 0.05 and 0.1 Hz in the respiration data (*T*_*sum*_ = 54.95, *p* = .015). However, this effect markedly excluded the spectral peak indicating that the magnitude of breathing remained constant over time (Fig. 2C). Finally, no significant changes were registered in the coupling between the two measures in each individual time window (Fig. 2D). These results also suggest that the negative power-power correlation reported above is likely emerging from a source of variability that is not explained by time passing (see e.g. Benwell et al., 2018 for time on task as a mediator in a different context), as breathing depth does not seem to change monotonically (decrease) across the three time windows.

Note that this effect is unlikely to be driven by physical changes in the environment during the task-related stimulation as this would likely produce a stronger spectral component in the pupil signal at the rate of the stimulus delivery (Schwiedrzik & Sudmann, 2020).

### Respiration-pupil coupling follows breathing rate

We explicitly tested the hypothesised influence of breathing depth on respiration-pupil coupling in a separate data set (N = 28, 14 female; Kluger et al., 2023). Participants were instructed to rest while breathing normally vs deeply in two separate 5-min measurements. Looking at the power spectra of pupil size and respiration time series, deeper breathing led to an overall decrease in respiratory rate, from the typical 0.27 Hz (*SD* = 0.06) to 0.21 Hz (*SD* = 0.07) on average (Wilcoxon signed rank test, *Z* = 3.60, *p* < .001; see Fig. 3A). Interestingly, pupillary dynamics did not exhibit a strong Hippus in this dataset. The typical peak (*M* = 0.26 Hz, *SD* = 0.13) seemed more pronounced during normal breathing than in the deep breathing condition (*M* = 0.24 Hz, *SD* = 0.12), however without producing measurable power differences between both conditions. Extracting Hippus peak frequencies therefore required removing the 1/f trend in the spectra (explained in detail in the Methods). Peak frequencies did not differ between breathing conditions (Wilcoxon signed rank test; *Z* = .697, *p* = .486; see Fig. 3B). Finally, we also observed substantial coupling between pupil and respiration in both conditions (normal breathing: *Tsum* = 14.75, *pcbpt* = .005; deep breathing: *Tsum* = 19.46, *pcbpt* = .007; also see Fig. 3C). Importantly, the peak frequency of the coupling also showed a substantial decrease from 0.26 (*SD* = 0.23) to 0.17 Hz (*SD* = 0.03), suggesting that the momentary respiratory rate strongly influences the coupling (Wilcoxon signed rank test; *Z* = 3.77, *p* < .001).

### Respiration-pupil coupling reduced during task performance

For a subset of the resting state data (N = 25) we were able to compare the above findings with similar data recorded while participants were performing a visual spatial detection task (Kluger et al., 2021). As quiescence and task performance have been shown to involve different levels of arousal (Podvalny et al., 2021; Reimer et al., 2014), we tested for a difference in coupling between respiration and pupil dynamics. Specifically, we compared the last of six blocks of task performance in an MEG experiment (length = 446 ± 34s [M ± SD]) with the immediately following resting state recording. We found evidence for decreased power of respiration during rest in frequency ranges above (0.38 - 0.52 Hz, *T*_*sum*_ = -6.81, *p*_*cbpt*_ = .044; 0.58 - 0.80 Hz, *T*_*sum*_ = -6.80, *p*_*cbpt*_ = .044; 1.12 - 6 Hz, *T*_*sum*_ = -42.51, *p*_*cbpt*_ < .001), but not including the respiratory peak frequency (see Fig. 3D). These differences may point at changes in oscillatory waveform of the respiratory rhythm that do not impact power at the fundamental frequency but only at higher harmonics (an index of non-sinusoidal properties of the time series). For the pupil, we observed systematically higher power in the Hippus frequency range (0.19 - 0.28 Hz; Fig. 3E) during rest (*T*_*sum*_ = 9.96, *p*_*cbpt*_ = .033), as well as a difference in the range between 0.90 - 2.46 Hz (*T*_*sum*_ = 11.11, *p*_*cbpt*_ = .026). This difference, and the fact that the Hippus is barely visible in the task data (Fig. 3E), are in line with earlier reports that an increased Hippus indicates decreased alertness (Mathot, 2018; McLaren, 1992). Ultimately, we contrasted the coupling in both conditions and found that coupling was stronger during rest within 0.16 to 0.31 frequency range (*T*_*sum*_ = 27.17, *p*_*cbpt*_ = .001; Fig. 3F), i.e., largely commensurate with the frequency range we observed the coupling in the resting state analysis above. Note that the weak Hippus in the task condition may fall within a low signal-to-noise regime where phase estimates will be affected by other signal components, and will therefore be less precise, which could contribute to a reduction in the coupling measure.

### Control - coupling across task blocks

To rule out that order effects contributed to task vs rest differences – the last task block was always presented prior to the rest block – we tested coupling across all six task blocks. Using the approaches outlined in the Methods section we tested for time on task-trends across the experimental blocks of Kluger and colleagues (2021), during which participants had to perform a task. We found a significant increase for frequencies below the peak frequency in the respiration data (Fig. 4A). For the pupil, cluster-based permutation testing returned one cluster including the Hippus frequency that showed a decrease over time (Fig. 4B). Importantly, no changes were found in coupling across blocks making it very unlikely that such trend would account for the task-rest differences (Fig. 4C).

**Figure 4.**
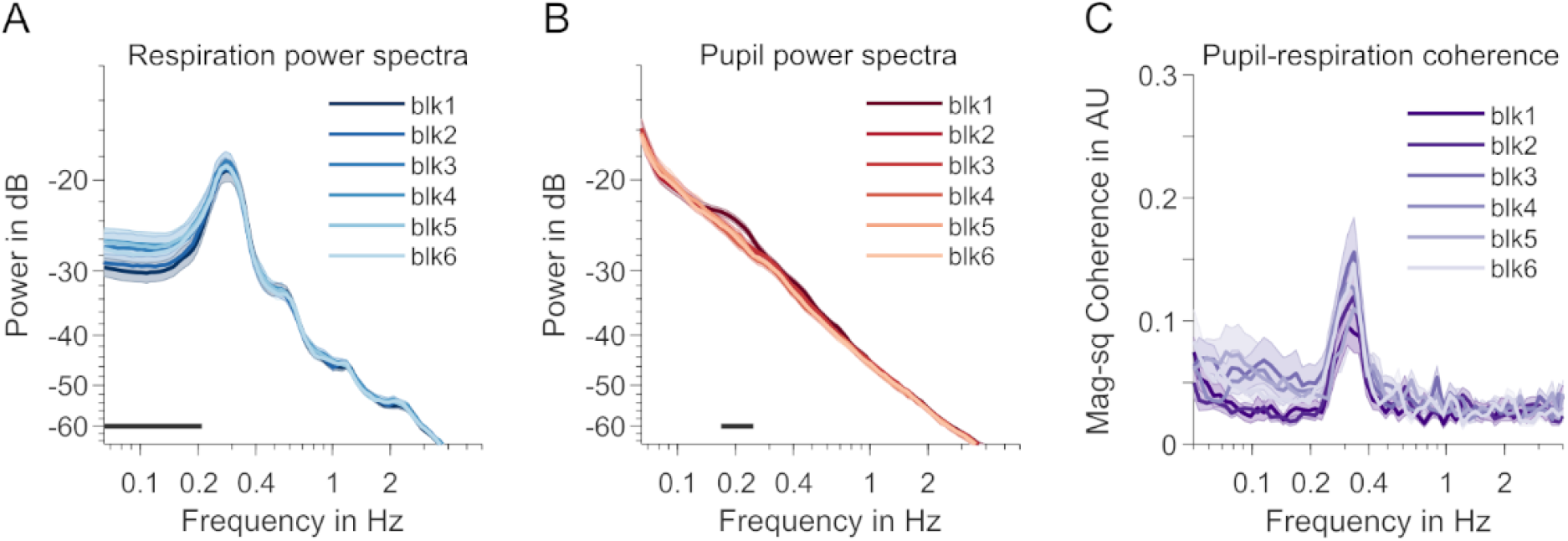
Differences between task blocks. **A** Respiration-power spectra across the six blocks including task performance. Light colours indicate later blocks. Shaded areas indicate SEM. A straight black line above the x-axis indicates a frequency cluster with a significant time-on-task effect (linear regression, p < .05, cluster-based permutation test). **B** Same as A but for pupil data. **C** Same as A & B but for respiration-pupil coupling (magnitude-squared coherence). No significant differences in coherence between blocks.

### Control - effect of pupil preprocessing

A concern in our analysis was whether our results could be influenced by residual artifacts in the pupil data, e.g., remainders of interpolated blinks. The spectral analysis, carried out on raw pupil time series below demonstrates that artifacts generally fall into a frequency range that is above the typical Hippus frequency (Fig. 5A), while producing a reduced, yet still above-chance measure of pupil-respiration coupling (Fig. 5B), thereby effectively ruling out an influence of basing our pupil analyses on blink-removed and interpolated time series, or of ocular artifacts on the present findings in general.

**Figure 5.**
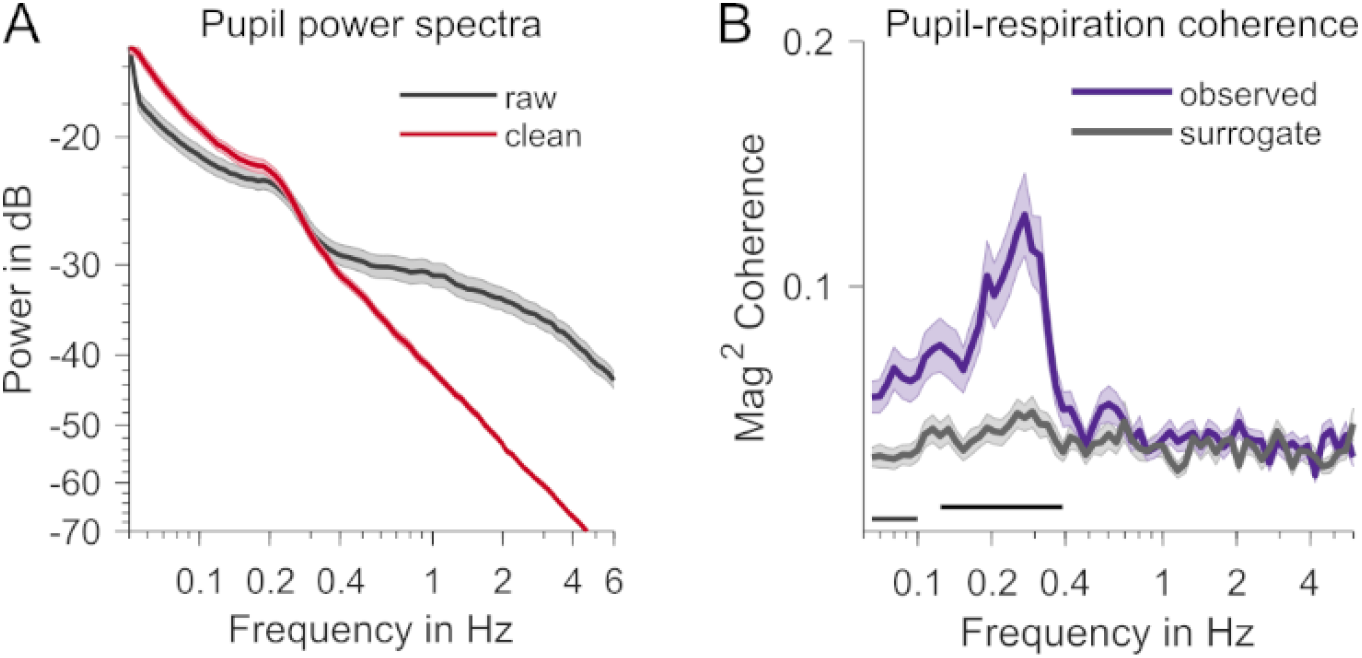
Effect of pupil preprocessing. **A** Pupil-power spectra during rest before (black) and after (red) preprocessing. **B** Coupling (magnitude-squared coherence) computed using raw, unprocessed pupil data. The purple trace shows the coupling observed in the data, the grey trace is based on surrogate data. Dark straight lines show frequency ranges with significant differences observed vs surrogate (p < .05, cluster-based permutation test).

## Discussion

In this investigation of respiration-arousal interplay, we demonstrate substantial phase coupling (magnitude-squared coherence) of pupil dilation dynamics to the breathing rhythm during rest. In an independent data set, changing the respiratory pattern towards voluntary deep breathing substantially decreased the peak frequency of respiration-pupil coupling. Finally, coupling decreased during a visual perception task (compared to rest) in a third data set, which opens up instructive avenues for comparing effects of contextual and respiratory interventions on phase coupling between respiration and pupil-linked arousal.

### Strong evidence for coupled rhythmic pupillary and respiratory dynamics

In their recent review of the literature on links between respiration and pupil dynamics, Schaefer and colleagues (Schaefer et al., 2023) graded the overall strength of evidence for the effect of breathing phase on pupil dynamics as “low”, and for the effects of breathing depth and rate as “very low”. Notably, their sample included studies with various measures of pupil dynamics (e.g., tonic pupil size, light reflex) and their correlations with respiration parameters. Focussing on oscillatory coupling in our investigation, we identified the rhythmic Hippus as a measure of interest, due to its prominent peak at around 0.2 Hz in spectra of pupil time series but also due its potential role as an indicator of momentary brain state (McLaren et al., 1992). More specifically, quantifying oscillatory coupling by means of phase coherence (e.g. Gross et al., 2021), an approach that had not been used before (see Schaefer et al., 2023), reliably produced a peak coherence between 0.2 - 0.3 Hz in three datasets (during normal breathing). Given that this is the typical range of the respiratory rate (Fleming et al., 2011), as well as the emerging Hippus during rest (Bouma and Baghuis, 1971; Turnbull et al., 2017), we provide strong evidence in favour of an effect of breathing phase (sensu Schaefer et al., 2023) on the Hippus aspect of pupillary dynamics (also see Schaefer et al., 2024 for the group’s latest preprint with similar results).

Given common reservations regarding directed functional connectivity (see above), our approach does not strictly allow a direct inference of whether respiration is the driver of the periodic arousal modulation linked to the Hippus. However, given the relative strength and constancy of the respiratory rhythm, a respiratory drive seems likely, and aligns with earlier ideas about the neural circuitry of this link (Borgdorff, 1975; Ohtsuka et al., 1988; Melnychuk et al., 2021) as well as its demonstration in the animal model (Yackle et al., 2017). It is therefore possible that the rhythmic Hippus can be explained by the entrainment of locus coeruleus neurons by the respiratory pacemaker in the pre-Bötzinger complex, a phenomenon known to occur between distinct but connected neuronal populations with the ability to produce concerted rhythmic activity (Thut et al., 2011; Herrmann et al., 2016).

Phase entrainment can explain the coupling, yet additional mechanisms need to be considered to explain the increase in Hippus magnitude at the peak frequency over the course of a 5 min resting state recording. Although this effect aligns with the idea that an increased Hippus indicates decreased vigilance or alertness as relaxation sets in (McLaren et al., 1992), it cannot simply be attributed to a similar increase in the magnitude of the fundamental respiratory rate, or a change in coupling strength because we found these to remain constant. Moreover, our results suggest that peak-frequency respiratory and Hippus magnitude may share an inverse relationship during rest, whereas an account in terms of entrainment (or resonance) would predict that increased magnitude in respiration leads to an increased Hippus. A more comprehensive understanding of the observed effects requires exploring factors that affected coupling. One could speculate that a third physiological influence, e.g. ultraslow gastric rhythms (0.05 Hz; Rebollo & Tallon-Baudry, 2022), other infraslow oscillations (Watson, 2018), or metastability of two or more physiological states (Kelso 2012) may mediate an anti-phasic relationship, where transient increases in the metabolic demand of oxygen indicate episodes of higher alertness as indicated by a reduced Hippus (and vice versa).

### Breathing mode and behavioural state alter pupil-respiration coupling

Changes we observed in pupil-respiration coupling based on manipulations of breathing mode (normal vs deep breathing) and behavioural state (rest vs task) shed further light on the interplay between respiratory and Hippus rhythms. Absent a direct measure of respiration, Pomè and colleagues (2020) have recently argued that observed changes in pupillary dynamics during mindfulness meditation may have been a consequence of altered breathing patterns. However, evidence for pupil size variations as a function of breathing depth has remained inconclusive (see Schaefer et al., 2023 and, to date, only few studies have contrasted the effects of different breathing modes directly. Debnath and colleagues (2021), for instance, investigated relationships between several physiological measures during a variety of physical exercises and stress tests that affected breathing and reported no overall effect. In contrast, Schumann and colleagues (2020) do report an increase in pupillary unrest during deep breathing, a measure for which they aggregate spectral power below 1.5 Hz. Similar to Schumann et al., we observed a marked decrease in breathing rate when participants were instructed to breathe deeply. Along with the change in breathing frequency, both the peak frequency of respiratory and Hippus signals decreased, as did the peak frequency of pupil-respiration coupling. This strong dependence of coupling frequency on breathing rate adds further support to the idea that the observed pupil-respiration link is primarily driven by the respiratory pacemaker. It may also provide an explanation for the inverse relationship between peak respiratory and Hippus power observed in the resting state data: Normal fluctuations in respiratory rate (peak frequency) between deeper and shallower (i.e., slower and faster) breathing lead to varying resonance in the neuromodulatory circuitry that influences pupil size.

In contrast to the frequency-changing effect of breathing mode, we observed a frequency-specific reduction in peak coupling strength (at around 0.2 Hz) when participants engaged in a task vs resting. Notably, this effect occurred in the absence of any peak frequency changes in respiration itself (although power differences in harmonic frequencies may point at waveform changes during task engagement), while the pupillary Hippus was effectively abolished during task performance. Together with its gradual increase during rest, this effect underlines its potential role as an index of current alertness, momentaneous internal vs external focus, and more generally, arousal levels where a higher Hippus indicates a state of lower arousal (Bouma and Baghuis, 1971; Mathôt, 2018; McLaren et al., 1992; Bouma and Baghuis, 1971; Mathôt, 2018). Taken together, the malleability of this indicator to differences in breathing mode suggest a strong respiratory drive, and the decrease in respiratory-coupling during task performance suggests that the respiratory drive may be more permeable in states of low arousal.

### Pupil-respiration coupling suggests interplay of breathing and arousal

Physiological arousal is primarily controlled via the noradrenergic neurotransmission as regulated by the locus coeruleus (LC; (Aston-Jones and Cohen, 2005; Mather and Harley, 2016). As the primary source of noradrenergic release in the mammalian brain, the LC projects to subcortical and cortical regions (Aston-Jones and Cohen, 2005). In concert with the release of acetylcholine (ACh) in the basal forebrain, the LC-NE axis profoundly influences cortical activity (Pfeffer et al., 2022; Podvalny et al., 2021; Radetz and Siegel, 2022) with consequences for cognitive function (Waschke et al., 2019; Kosciessa et al., 2021). An intriguing possibility then is that respiratory rhythms constitute another, potentially earlier node in a sequence of physiological processes that modulate cortical activity by controlling noradrenaline release, creating a causal loop between interoceptive sensation, respiration, and arousal. In line with this, a recent study by Yackle and colleagues (2017) identified a subpopulation of neurons in the preBötzinger Complex, the primary respiratory rhythm generator of the brainstem, with direct projections to noradrenaline expressing LC neurons. Ablating these connections further eliminated the breath–by–breath control of noradrenaline release, causing mice to exhibit altered arousal responses to exteroceptive stimuli.

In the light of these connections in the underlying neural and neuromodulatory circuitry, and our finding that pupil-respiratory coupling is stronger during a restful, introspective behavioural state, we can speculate that the brain may adopt a metabolically optimal resting mode that is characterised by a rhythmic intermittent release of neuromodulators, instead of a tonic, more continuous release during active performance that requires a higher sustained level of cortical activation for the processing of task-relevant sensory input (see Schroeder and Lakatos, 2009 for a similar suggestion regarding the functional significance of cortical electrophysiological oscillations). This account allows testable predictions: Cortical markers of sensory processing may underlie stronger periodic fluctuations with frequencies in the range of the coupling observed here when participants are in a more restful than alert state.

In summary, we show evidence for an oscillatory coupling of respiratory and pupil dynamics that critically depends on breathing phase and rate, and also changes depending on behavioural state. These results contrast with the results of a recent survey which showed low confidence overall in pupil-respiration links (Schaefer et al., 2023). However, the survey considered a range of possible links, whereas we focussed on oscillatory coupling, possibly explaining the divergence, and pointing out a promising direction to take to explore these links further. The close link between pupil dynamics and neuromodulatory influences of the arousal system on cortical function on one side, and the potential drive of pupil-linked arousal by respiration, poses new exciting challenges for our understanding of how physiological processes influence cortical activity, and hence, human cognition.

## Acknowledgments

The authors would like to thank Karin Wilken, Ute Trompeter, and Hildegard Deitermann for their help with data collection.

## Funding

DSK is supported by the DFG (grant number KL 3580/1-1) and the IMF (KL 1 2 22 01). JG is supported by the DFG (GR 2024/11-1, GR 2024/12-1). We acknowledge support from the Open Access Publication Fund of the University of Münster. All authors are members of the Scottish-EU Critical Oscillations Network (SCONe), funded by the Royal Society of Edinburgh (RSE Saltire Facilitation Network Award to CK, Reference Number 1963).

## Author contributions

Contributions coded according to the CRediT taxonomy (Brand et al., 2015). Conceptualization: DSK, CK. Data curation: DSK, CK. Formal analysis: DSK, CK. Funding acquisition: DSK, JG. Investigation: DSK. Methodology: All authors. Project administration: CK, JG. Resources: DSK, JG. Software: DSK, CK. Supervision: CK, JG. Validation: CK, JG. Visualization: DSK, CK. Writing – original draft: DSK, CK. Writing – review & editing: All authors. Final version of the manuscript approved by all authors.

## Declaration of conflicting interests

The authors declare that there were no competing interests with respect to the authorship or publication of this research article.

## Notes

### Competing Interest Statement

The authors have declared no competing interest.

### Summary of Updates

Figs 1 and 3 revised; new directed connectivity analysis

## References

Al, E., Iliopoulos, F., Forschack, N., Nierhaus, T., Grund, M., Motyka, P., Gaebler, M., Nikulin, V.V., Villringer, A., 2020. Heart-brain interactions shape somatosensory perception and evoked potentials. Proc. Natl. Acad. Sci. USA 117, 10575–10584. doi:10.1073/pnas.1915629117

Allen, M., Poggiali, D., Whitaker, K., Marshall, T.R., Kievit, R.A., 2019. Raincloud plots: a multi-platform tool for robust data visualization. [version 1; peer review: 2 approved]. Wellcome Open Res. 4, 63. doi:10.12688/wellcomeopenres.15191.1

Allen, M., Varga, S., Heck, D.H., 2023. Respiratory rhythms of the predictive mind. Psychol. Rev. 130, 1066–1080. doi:10.1037/rev0000391

Aston-Jones, G., Cohen, J.D., 2005. An integrative theory of locus coeruleus-norepinephrine function: adaptive gain and optimal performance. Annu. Rev. Neurosci. 28, 403–450. doi:10.1146/annurev.neuro.28.061604.135709

Bastos, A. M., & Schoffelen, J. M. (2016). A tutorial review of functional connectivity analysis methods and their interpretational pitfalls. Frontiers in systems neuroscience, 9, 175.

Benwell, C.S.Y., Keitel, C., Harvey, M., Gross, J., Thut, G., 2018. Trial-by-trial co-variation of pre-stimulus EEG alpha power and visuospatial bias reflects a mixture of stochastic and deterministic effects. Eur. J. Neurosci. 48, 2566–2584. doi:10.1111/ejn.13688

Borgdorff, P., 1975. Respiratory fluctuations in pupil size. Am. J. Physiol. 228, 1094–1102. doi:10.1152/ajplegacy.1975.228.4.1094

Bouma, H., Baghuis, L.C., 1971. Hippus of the pupil: periods of slow oscillations of unknown origin. Vision Res. 11, 1345–1351. doi:10.1016/0042-6989(71)90016-2

Brændholt, M., Kluger, D.S., Varga, S., Heck, D.H., Gross, J., Allen, M.G., 2023. Breathing in waves: Understanding respiratory-brain coupling as a gradient of predictive oscillations. Neurosci. Biobehav. Rev. 152, 105262. doi:10.1016/j.neubiorev.2023.105262

Brand, A., Allen, L., Altman, M., Hlava, M., Scott, J., 2015. Beyond authorship: attribution, contribution, collaboration, and credit. Learn. Pub. 28, 151–155. doi:10.1087/20150211

Bressler, S. L., & Seth, A. K. (2011). Wiener–Granger causality: a well established methodology. Neuroimage, 58(2), 323–329.

Carter, G., Knapp, C., Nuttall, A., 1973. Estimation of the magnitude-squared coherence function via overlapped fast Fourier transform processing. IEEE Trans. Audio Electroacoust. 21, 337–344. doi:10.1109/TAU.1973.1162496

Criscuolo, A., Schwartze, M., Kotz, S.A., 2022. Cognition through the lens of a body-brain dynamic system. Trends Neurosci. doi:10.1016/j.tins.2022.06.004

Debnath, S., Levy, T.J., Bellehsen, M., Schwartz, R.M., Barnaby, D.P., Zanos, S., Volpe, B.T., Zanos, T.P., 2021. A method to quantify autonomic nervous system function in healthy, able-bodied individuals. Bioelectron. Med. 7, 13. doi:10.1186/s42234-021-00075-7

Del Negro, C.A., Funk, G.D., Feldman, J.L., 2018. Breathing matters. Nat. Rev. Neurosci. 19, 351–367. doi:10.1038/s41583-018-0003-6

Fleming, S., Thompson, M., Stevens, R., Heneghan, C., Plüddemann, A., Maconochie, I., Tarassenko, L., Mant, D., 2011. Normal ranges of heart rate and respiratory rate in children from birth to 18 years of age: a systematic review of observational studies. Lancet 377, 1011–1018. doi:10.1016/S0140-6736(10)62226-X

Galvez-Pol, A., McConnell, R., Kilner, J.M., 2020. Active sampling in visual search is coupled to the cardiac cycle. Cognition 196, 104149. doi:10.1016/j.cognition.2019.104149

Groot, J.M., Boayue, N.M., Csifcsák, G., Boekel, W., Huster, R., Forstmann, B.U., Mittner, M., 2021. Probing the neural signature of mind wandering with simultaneous fMRI-EEG and pupillometry. Neuroimage 224, 117412. doi:10.1016/j.neuroimage.2020.117412

Gross, J., Kluger, D.S., Abbasi, O., Chalas, N., Steingräber, N., Daube, C., Schoffelen, J.-M., 2021. Comparison of undirected frequency-domain connectivity measures for cerebro-peripheral analysis. Neuroimage 245, 118660. doi:10.1016/j.neuroimage.2021.118660

Harris, K.D., Thiele, A., 2011. Cortical state and attention. Nat. Rev. Neurosci. 12, 509–523. doi:10.1038/nrn3084

Hasselmo, M.E., 1995. Neuromodulation and cortical function: modeling the physiological basis of behavior. Behav. Brain Res. 67, 1–27. doi:10.1016/0166-4328(94)00113-t

Herrmann, C.S., Murray, M.M., Ionta, S., Hutt, A., Lefebvre, J., 2016. Shaping Intrinsic Neural Oscillations with Periodic Stimulation. J. Neurosci. 36, 5328–5337. doi:10.1523/JNEUROSCI.0236-16.2016

Hoeks, B., Levelt, W.J.M., 1993. Pupillary dilation as a measure of attention: a quantitative system analysis. Behavior Research Methods, Instruments, & Computers 25, 16–26. doi:10.3758/BF03204445

Kelso, J. S. (2012). Multistability and metastability: understanding dynamic coordination in the brain. Philosophical Transactions of the Royal Society B: Biological Sciences, 367(1591), 906–918.

Kluger, D.S., Gross, J., 2020. Depth and phase of respiration modulate cortico-muscular communication. Neuroimage 222, 117272. doi:10.1016/j.neuroimage.2020.117272

Kluger, D.S., Balestrieri, E., Busch, N.A., Gross, J., 2021. Respiration aligns perception with neural excitability. Elife 10. doi:10.7554/eLife.70907

Kluger, D.S., Forster, C., Abbasi, O., Chalas, N., Villringer, A., Gross, J., 2023. Modulatory dynamics of periodic and aperiodic activity in respiration-brain coupling. Nat. Commun. 14, 4699. doi:10.1038/s41467-023-40250-9

Kluger, D. S., Allen, M. G., & Gross, J., 2024. Brain–body states embody complex temporal dynamics. Trends in Cognitive Sciences. doi: 10.1016/j.tics.2024.05.003

Knapen, T., de Gee, J.W., Brascamp, J., Nuiten, S., Hoppenbrouwers, S., Theeuwes, J., 2016. Cognitive and Ocular Factors Jointly Determine Pupil Responses under Equiluminance. PLoS One 11, e0155574. doi:10.1371/journal.pone.0155574

Kosciessa, J.Q., Lindenberger, U., Garrett, D.D., 2021. Thalamocortical excitability modulation guides human perception under uncertainty. Nat. Commun. 12, 2430. doi:10.1038/s41467-021-22511-7

Lee, S.-H., Dan, Y., 2012. Neuromodulation of brain states. Neuron 76, 209–222. doi:10.1016/j.neuron.2012.09.012

Mather, M., Harley, C.W., 2016. The locus coeruleus: essential for maintaining cognitive function and the aging brain. Trends Cogn. Sci. (Regul. Ed.) 20, 214–226. doi:10.1016/j.tics.2016.01.001

Mathôt, S., 2018. Pupillometry: psychology, physiology, and function. J. Cogn. 1, 16. doi:10.5334/joc.18

McGinley, M.J., David, S.V., McCormick, D.A., 2015a. Cortical membrane potential signature of optimal states for sensory signal detection. Neuron 87, 179–192. doi: 10.1016/j.neuron.2015.05.038

McGinley, M.J., Vinck, M., Reimer, J., Batista-Brito, R., Zagha, E., Cadwell, C.R., Tolias, A.S., Cardin, J.A., McCormick, D.A., 2015b. Waking state: rapid variations modulate neural and behavioral responses. Neuron 87, 1143–1161. doi:10.1016/j.neuron.2015.09.012

McLaren, J.W., Erie, J.C., Brubaker, R.F., 1992. Computerized analysis of pupillograms in studies of alertness. Invest. Ophthalmol. Vis. Sci. 33, 671–676.

Melnychuk, M. C., Robertson, I. H., Plini, E. R., & Dockree, P. M., 2021. A bridge between the breath and the brain: synchronization of respiration, A pupillometric marker of the locus coeruleus, and an EEG marker of attentional control state. Brain sciences, 11(10), 1324. doi: 10.3390/brainsci11101324

Nakamura, N.H., Fukunaga, M., Oku, Y., 2019. Respiratory fluctuations in pupil diameter are not maintained during cognitive tasks. Respir. Physiol. Neurobiol. 265, 68–75. doi:10.1016/j.resp.2018.07.005

Ohtsuka, K., Asakura, K., Kawasaki, H., Sawa, M., 1988. Respiratory fluctuations of the human pupil. Exp. Brain Res. 71, 215–217. doi:10.1007/BF00247537

Oostenveld, R., Fries, P., Maris, E., Schoffelen, J.-M., 2011. FieldTrip: Open source software for advanced analysis of MEG, EEG, and invasive electrophysiological data. Comput Intell Neurosci 2011, 156869. doi:10.1155/2011/156869

Park, H.-D., Barnoud, C., Trang, H., Kannape, O.A., Schaller, K., Blanke, O., 2020. Breathing is coupled with voluntary action and the cortical readiness potential. Nat. Commun. 11, 289. doi:10.1038/s41467-019-13967-9

Pfeffer, T., Keitel, C., Kluger, D.S., Keitel, A., Russmann, A., Thut, G., Donner, T.H., Gross, J., 2022. Coupling of pupil- and neuronal population dynamics reveals diverse influences of arousal on cortical processing. Elife 11. doi:10.7554/eLife.71890

Podvalny, E., King, L.E., He, B.J., 2021. Spectral signature and behavioral consequence of spontaneous shifts of pupil-linked arousal in human. Elife 10. doi:10.7554/eLife.68265

Pomè, A., Burr, D.C., Capuozzo, A., Binda, P., 2020. Spontaneous pupillary oscillations increase during mindfulness meditation. Curr. Biol. 30, R1030–R1031. doi:10.1016/j.cub.2020.07.064

Radetz, A., Siegel, M., 2022. Spectral fingerprints of cortical neuromodulation. J. Neurosci. 42, 3836–3846. doi:10.1523/JNEUROSCI.1801-21.2022

Rebollo, I., Devauchelle, A.-D., Béranger, B., Tallon-Baudry, C., 2018. Stomach-brain synchrony reveals a novel, delayed-connectivity resting-state network in humans. Elife 7. doi:10.7554/eLife.33321

Rebollo, I., Wolpert, N., Tallon-Baudry, C., 2021. Brain–stomach coupling: Anatomy, functions, and future avenues of research. Curr. Opin. Biomed. Eng 18, 100270. doi:10.1016/j.cobme.2021.100270

Rebollo, I., & Tallon-Baudry, C. (2022). The sensory and motor components of the cortical hierarchy are coupled to the rhythm of the stomach during rest. Journal of Neuroscience, 42(11), 2205–2220.

Reimer, J., Froudarakis, E., Cadwell, C.R., Yatsenko, D., Denfield, G.H., Tolias, A.S., 2014. Pupil fluctuations track fast switching of cortical states during quiet wakefulness. Neuron 84, 355–362. doi:10.1016/j.neuron.2014.09.033

Schaefer, M., Edwards, S., Nordén, F., Lundström, J.N., Arshamian, A., 2023. Inconclusive evidence that breathing shapes pupil dynamics in humans: a systematic review. Pflugers Arch. 475, 119–137. doi:10.1007/s00424-022-02729-0

Schaefer, M., Mathot, S., Lundqvist, M., Lundstrom, J. N., & Arshamian, A. (2024). The Respiratory-Pupillary Phase Effect: Pupils size is smallest around inhalation onset and largest during exhalation. bioRxiv, 2024–06.

Schneider, M., Hathway, P., Leuchs, L., Sämann, P.G., Czisch, M., Spoormaker, V.I., 2016. Spontaneous pupil dilations during the resting state are associated with activation of the salience network. Neuroimage 139, 189–201. doi:10.1016/j.neuroimage.2016.06.011

Schreiber, T., Schmitz, A., 1996. Improved surrogate data for nonlinearity tests. Phys. Rev. Lett. 77, 635–638. doi:10.1103/PhysRevLett.77.635

Schroeder, C.E., Lakatos, P., 2009. Low-frequency neuronal oscillations as instruments of sensory selection. Trends Neurosci. 32, 9–18. doi:10.1016/j.tins.2008.09.012

Schumann, A., Kietzer, S., Ebel, J., Bär, K.J., 2020. Sympathetic and parasympathetic modulation of pupillary unrest. Front. Neurosci. 14, 178. doi:10.3389/fnins.2020.00178

Schwalm, M., Rosales Jubal, E., 2017. Back to pupillometry: how cortical network state fluctuations tracked by pupil dynamics could explain neural signal variability in human cognitive neuroscience. eNeuro 4. doi:10.1523/ENEURO.0293-16.2017

Schwiedrzik, C. M., & Sudmann, S. S. (2020). Pupil diameter tracks statistical structure in the environment to increase visual sensitivity. Journal of Neuroscience, 40(23), 4565–4575.

Smulders, F.T.Y., 2010. Simplifying jackknifing of ERPs and getting more out of it: retrieving estimates of participants’ latencies. Psychophysiology 47, 387–392. doi:10.1111/j.1469-8986.2009.00934.x

Steriade, M., 1996. Arousal: revisiting the reticular activating system. Science 272, 225–226. doi:10.1126/science.272.5259.225

Thut, G., Schyns, P.G., Gross, J., 2011. Entrainment of perceptually relevant brain oscillations by non-invasive rhythmic stimulation of the human brain. Front. Psychol. 2, 170. doi:10.3389/fpsyg.2011.00170

Turnbull, P.R.K., Irani, N., Lim, N., Phillips, J.R., 2017. Origins of pupillary hippus in the autonomic nervous system. Invest. Ophthalmol. Vis. Sci. 58, 197–203. doi:10.1167/iovs.16-20785

Urai, A.E., Braun, A., Donner, T.H., 2017. Pupil-linked arousal is driven by decision uncertainty and alters serial choice bias. Nat. Commun. 8, 14637. doi:10.1038/ncomms14637

Vinck, M., Batista-Brito, R., Knoblich, U., Cardin, J.A., 2015. Arousal and locomotion make distinct contributions to cortical activity patterns and visual encoding. Neuron 86, 740–754. doi:10.1016/j.neuron.2015.03.028

Waschke, L., Tune, S., Obleser, J., 2019. Local cortical desynchronization and pupil-linked arousal differentially shape brain states for optimal sensory performance. Elife 8. doi:10.7554/eLife.51501

Watson, B. O. (2018). Cognitive and physiologic impacts of the infraslow oscillation. Frontiers in systems neuroscience, 12, 44.

Wierda, S.M., van Rijn, H., Taatgen, N.A., Martens, S., 2012. Pupil dilation deconvolution reveals the dynamics of attention at high temporal resolution. Proc. Natl. Acad. Sci. USA 109, 8456–8460. doi:10.1073/pnas.1201858109

Yackle, K., Schwarz, L.A., Kam, K., Sorokin, J.M., Huguenard, J.R., Feldman, J.L., Luo, L., Krasnow, M.A., 2017. Breathing control center neurons that promote arousal in mice. Science 355, 1411–1415. doi:10.1126/science.aai7984

Yang, C.F., Feldman, J.L., 2018. Efferent projections of excitatory and inhibitory preBötzinger Complex neurons. J. Comp. Neurol. 526, 1389–1402. doi:10.1002/cne.24415

